# Exploratory Multi-Omics Profiling of Rheumatoid Arthritis Heterogeneity

**DOI:** 10.1101/2025.11.17.688813

**Authors:** Niket Rajeevan, Zain Khan

## Abstract

Rheumatoid arthritis (RA) is a clinically heterogeneous disease with highly variable treatment responses despite widespread use of conventional and biologic disease-modifying therapies. Existing clinical and serological markers only partially explain this variability and are not routinely used to personalise drug choice. We hypothesised that integrated plasma proteomic– metabolomic profiling could capture dimensions of RA molecular heterogeneity that are invisible to standard phenotyping and highlight characteristic patterns of metabolic dysregulation.

We profiled 384 proteins and 1,063 metabolites in 80 RA patients and 40 healthy controls. Unsupervised consensus clustering of the integrated omics matrix identified two stable molecular subtypes within the RA cohort. Anti-citrullinated protein antibody (ACPA) status, which in this cohort was evenly split overall (40 ACPA-positive, 40 ACPA-negative), was similarly distributed across the two subtypes, suggesting that the molecular groupings reflect heterogeneity orthogonal to conventional serology. Protein–metabolite association network analysis revealed a substantial reduction in association edges in RA versus controls, with pathway-specific effects: pronounced connectivity loss in amino-acid metabolism but paradoxical gains in inflammatory mediators including glutamate and sphingolipids. Preserved correlation strength among surviving edges is consistent with selective pathway disconnection rather than uniformly weakened coordination.

These hypothesis-generating findings, limited by cross-sectional design, medication heterogeneity, and single-centre recruitment, suggest that RA comprises molecularly distinct subtypes not captured by standard serological categories and is associated with extensive metabolic network reorganisation. This multi-omics framework illustrates how network-level signatures could support future precision stratification efforts and motivates further investigation of metabolic network restoration as a potential therapeutic avenue.

## 1 Introduction

Rheumatoid arthritis (RA) is a chronic autoimmune inflammatory disease affecting millions of people worldwide and remains associated with substantial morbidity despite modern treat-to-target strategies. Treatment outcomes are heterogeneous: many patients fail to achieve sustained remission or low disease activity even after sequential biologic or targeted synthetic DMARDs [1, 2].

One way RA is commonly described is as anti-citrullinated protein antibody (ACPA)-positive or ACPA-negative [3]. However, conventional serology captures only part of the clinically relevant RA heterogeneity. Marked variation in disease course, treatment response, and extra-articular manifestations implies underlying molecular diversity that is not fully explained by autoantibody status alone [4, 5]. Synovial-tissue transcriptomic studies have identified discrete cellular and molecular “pathotypes” that correlate with biologic and conventional synthetic DMARD response, supporting the existence of biologically distinct RA subsets [4, 5]. In parallel, plasma and serum metabolomics studies highlight widespread alterations in amino-acid, lipid, and energy metabolism linked to disease activity and clinical complications [6].

Most of these studies interrogate a single molecular layer at a time. Multi-omics approaches integrating genomics, transcriptomics, proteomics, metabolomics, and the microbiome are now beginning to reveal coordinated alterations across modalities and to support refined molecular patient stratification in RA [7, 8]. Integrated proteomic–metabolomic profiling in particular offers a pragmatic route to capture both inflammatory mediators and downstream metabolic consequences within a single plasma sample.

We therefore performed exploratory plasma proteomic and metabolomic profiling in 80 RA patients (40 ACPA-positive, 40 ACPA-negative) and 40 healthy controls. Our objectives were twofold: first, to determine whether molecular subtypes exist that are not aligned with ACPA classification; and second, to characterise protein–metabolite association networks in RA to test whether the disease is associated with selective metabolic rewiring or more diffuse dysregulation. These hypothesis-generating analyses aim to provide proof-of-concept for multi-omics-based patient stratification in RA.

### 1.1 Study Objectives

We addressed two complementary objectives. First, we applied unsupervised clustering to integrated proteomic–metabolomic data to test whether molecularly distinct patient subgroups exist within RA beyond conventional ACPA classification (**Objective 1**). If robust molecular endotypes emerge independent of serology, this would support stratified therapeutic approaches [4, 5].

Second, we characterized protein–metabolite association networks to examine how disease disrupts metabolic coordination (**Objective 2**). Healthy cells maintain metabolic homeostasis through coordinated protein–metabolite relationships; in disease, these association patterns may show systematic disruption [9, 10]. Network analysis—examining correlation patterns between proteins and metabolites—can distinguish random dysregulation from selective pathway rewiring with therapeutic implications [11].

These objectives are synergistic: molecular subgrouping identifies *which* patients differ, while network analysis reveals *how* their metabolic coordination differs. If distinct subgroups exhibit different association network disruption patterns, this would suggest that therapeutic strategies should target subtype-specific metabolic vulnerabilities rather than inflammation uniformly [12, 13].

### 1.2 Dataset and Study Population

We analyzed plasma multi-omics data [14] from 120 individuals: 40 ACPA-negative RA patients, 40 ACPA-positive RA patients, and 40 healthy controls. All RA patients met 2010 ACR/EULAR classification criteria [15] and exhibited heterogeneous disease activity (DAS28-CRP range 1.5–7.5, mean 3.5). Patients were on diverse therapeutic regimens including methotrexate, other conventional synthetic DMARDs, prednisone, and biologics – representing an important potential confounder in molecular analyses. Groups were matched for age, sex, BMI, and smoking history.

Proteomic profiling used an aptamer-based platform (SomaScan) [16] quantifying 7,273 proteins, with 384 retained after quality control. Metabolomic profiling employed UPLC-MS/MS, yielding 1,063 metabolites after removing features with *>*20% missing values and applying normalization procedures. Clinical metadata included DAS28-CRP, medication history, and autoantibody status.

## 2 Results and Discussion

### 2.1 Molecular Subgroup Discovery

To test whether RA patients segregate into molecularly distinct subgroups beyond conventional ACPA classification, we employed consensus clustering–a rigorous resampling approach that identifies only patient groupings stable across multiple analytical variations. Single clustering runs can produce spurious groupings dependent on specific algorithmic choices or random initializations. Consensus clustering addresses this by performing 1000 iterations of hierarchical clustering, each randomly subsampling 80% of patients and 90% of features while varying the clustering algorithm (Ward’s, average, and complete linkage methods). For each iteration, we recorded which patient pairs clustered together. The final consensus score represents the overall stability of cluster assignments across all iterations–effectively a measure of clustering confidence independent of any single algorithmic choice [17, 18].

This analysis identified two highly stable molecular subtypes with an overall consensus score of 0.898, indicating strong reproducibility across all analytical perturbations. Critically, these molecular subtypes were statistically independent of ACPA serological status (*χ*^2^ p=0.789). The larger subtype (Cluster 1, n=62) comprised 32 ACPA-positive and 30 ACPA-negative patients; the smaller subtype (Cluster 2, n=18) contained 8 ACPA-positive and 10 ACPA-negative patients. This near-equal distribution across both subtypes demonstrates that comprehensive molecular profiling captures heterogeneity orthogonal to autoantibody testing–patients with and without ACPA are represented in both molecular subtypes [4, 5].

Visualization in principal component space (Figure 1A) confirms the separation, with Cluster 2 forming a more compact, distinct group along the first principal component, which captures 28.6% of molecular variance. Patient-level assignment stability was remarkably high: 72 of 80 patients (90%) exhibited greater than 80% consensus across iterations, meaning they clustered with the same group of patients in over 80 of the 100 iterations. The distribution of individual stability scores (Figure 1B) shows a bimodal pattern heavily skewed toward high stability (mean=0.879), indicating most patients have unambiguous subtype membership. This distribution–with a sharp peak at high stability and few intermediate-stability patients–suggests differing molecular states. Differential expression analysis identified 254 proteins and metabolites significantly discriminating the two subtypes (FDR *<* 0.001, Mann-Whitney U test). Top discriminators included metabolic enzymes (glutaminase [GLS], glucose-6-phosphate dehydrogenase [G6PD]), inflammatory mediators (CXCL3, phospholipase A2 [PLA2G4A]), and PI3K/AKT pathway components (AKT1, PDPK1), consistent with coordinated immunometabolic reprogramming distinguishing the subtypes.

**Figure 1.**
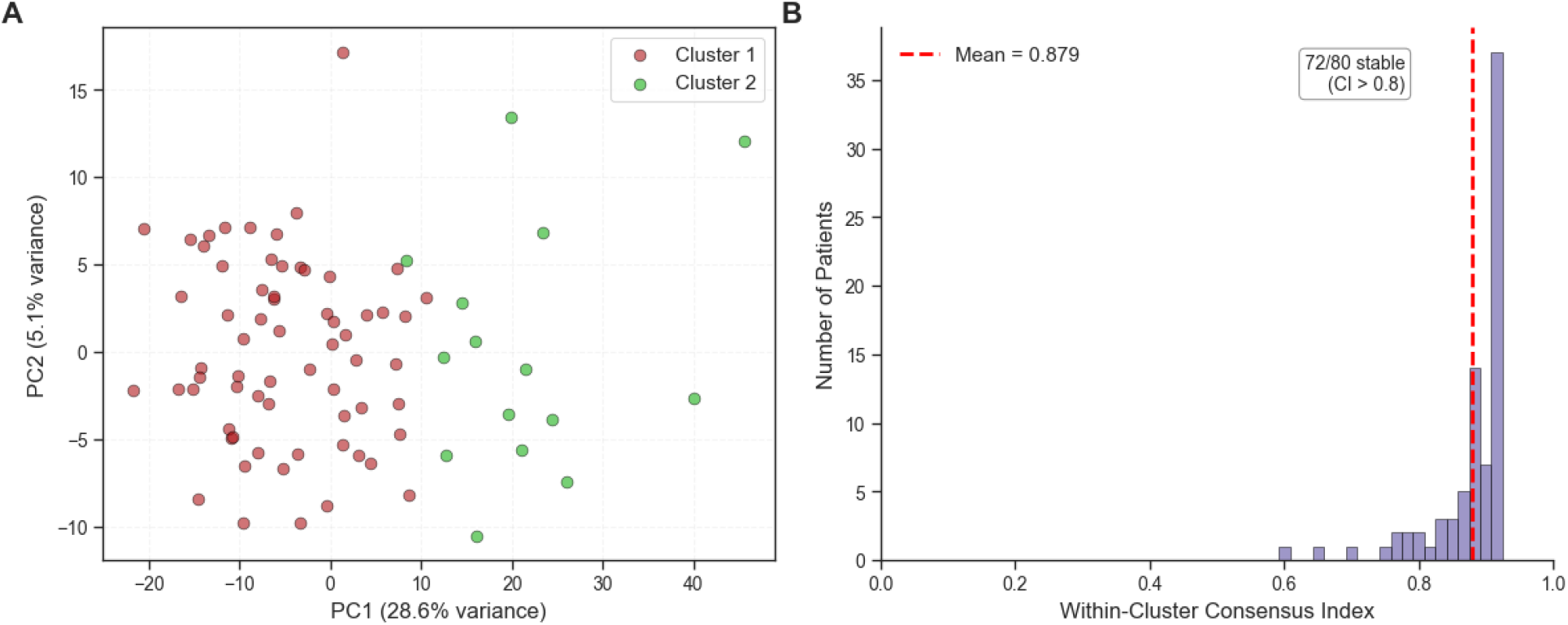
Consensus clustering reveals two stable molecular subtypes independent of ACPA status. (A) Consensus clusters visualized in PCA space show clear separation along PC1 (28.6% variance), with Cluster 2 (orange) forming a compact, distinct molecular subtype. (B) Distribution of individual patient stability scores shows bimodal pattern with mean=0.879, indicating robust, discrete molecular states with 72/80 patients exhibiting *>*80% consensus.

These exploratory findings support the hypothesis that integrated multi-omics profiling captures molecular heterogeneity not evident from ACPA testing alone [5]. Current RA classification relies primarily on a single serological marker, yet our data suggest that comprehensive molecular profiling reveals additional axes of biological variation. However, external validation in independent cohorts is required to confirm subtype robustness, assess longitudinal stability, and establish whether these molecular distinctions have clinical utility for treatment stratification.

### 2.2 Protein–Metabolite Association Network Analysis

Having identified molecular subtypes, we next examined how disease disrupts metabolic coordination at the systems level. In healthy cells, proteins (primarily enzymes) and metabolites interact in coordinated networks: enzymes regulate metabolite levels, which in turn provide feedback to protein activity. These association networks help maintain metabolic homeostasis. In disease, these coordination patterns may become disrupted in systematic ways that reveal underlying pathological mechanisms [9, 10].

We constructed protein–metabolite association networks for RA and control groups using Spearman correlations, defining network “edges” (connections) by correlation strength (|*r*| *>* 0.4) and statistical significance (*p <* 0.01, uncorrected). While these thresholds are exploratory and results are threshold-sensitive, this framework provides initial hypothesis-generating evidence for how metabolic coordination may differ between health and disease [9].

Network analysis revealed a substantial reduction in association density in RA versus controls (58% fewer edges: 9,057 vs. 21,571; permutation *p <* 0.001). The network density heatmap (Figure 2A) visualises this disruption: metabolites showing robust connectivity in controls (red bands) exhibit predominantly sparse connectivity in RA (blue). This pattern is consistent with widespread loss of coordinated metabolic regulation, although it likely reflects a composite of disease-intrinsic biology, medication effects, and inflammatory activity rather than intrinsic disease mechanisms alone [19, 20].

**Figure 2.**
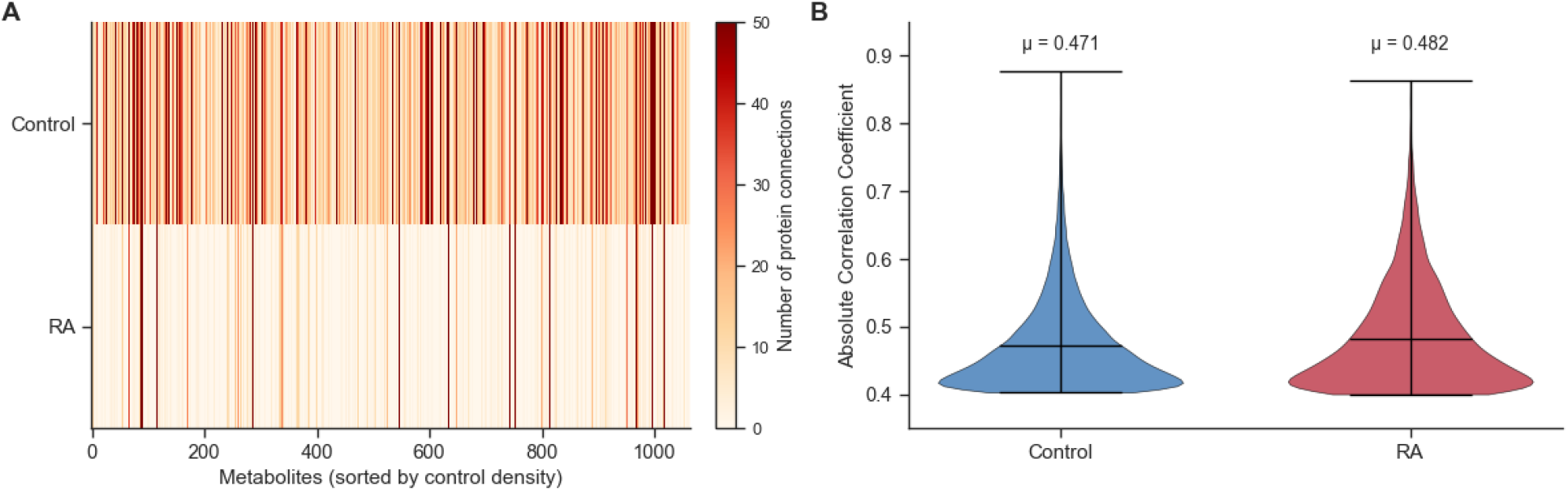
Protein-metabolite association networks show selective disruption in RA. (A) Network density heatmap (metabolites sorted by control connectivity) reveals reduced RA connectivity patterns. Red indicates high connectivity; blue/white indicates sparse connectivity. (B) Preserved correlation strength despite edge loss suggests selective pathway disconnection rather than uniformly weakened regulation.

Importantly, despite this large reduction in the number of edges, the correlation strength of remaining connections was preserved (control mean |*r*| = 0.471 vs. RA mean |*r*| = 0.482; Figure 2B). This observation suggests that RA is not characterised by a generalised weakening of all metabolic relationships but rather by selective disconnection of specific pathways while others remain strongly coordinated—a pattern more consistent with active metabolic reprogramming than with purely random degradation [20, 21].

The volcano plot (Figure 3) highlights pathway specificity in this reorganisation. Most metabolites lost connectivity (red points to the left), particularly those involved in amino-acid metabolism, consistent with disrupted homeostatic metabolic wiring. However, a subset of metabolites showed paradoxical connectivity gains. Glutamate (+139 edges), aspartate (+101 edges), and sphingolipids (+99 edges) exhibited increased associations despite the overall network contraction. These molecules have established roles in inflammatory biology: glutamine/glutamate metabolism supports effector T cells, macrophages, and fibroblast-like synoviocytes in RA and other immunemediated diseases [19, 20, 23]; aspartate is a key precursor for de novo nucleotide biosynthesis and is indispensable for proliferating cells [22]; and sphingolipid metabolites such as ceramide and sphingosine-1-phosphate regulate inflammatory signalling and immune cell trafficking [24].

**Figure 3.**
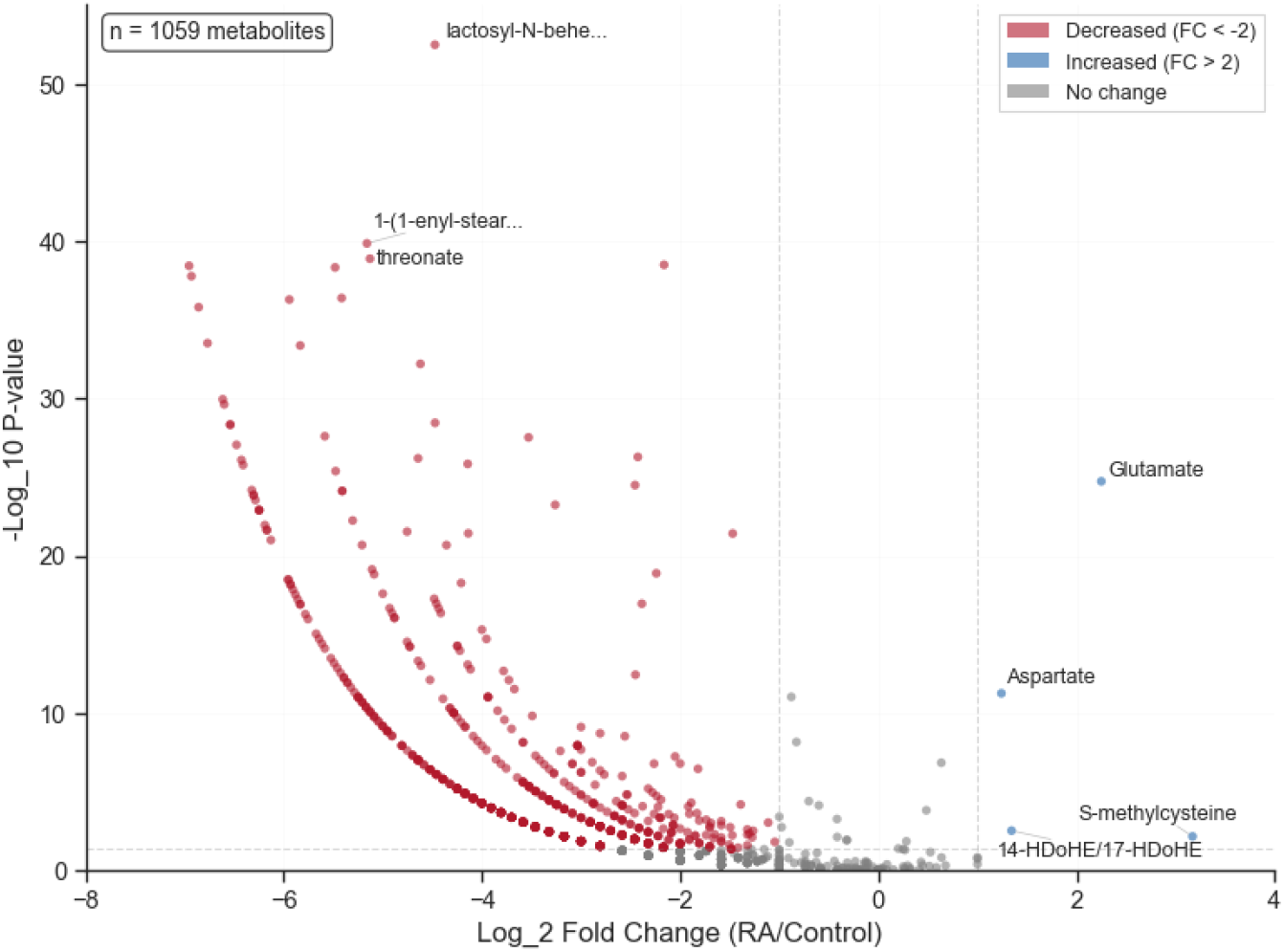
Volcano plot quantifies metabolite connectivity changes: amino acid metabolism losses (red, left) versus inflammatory metabolite gains (blue, right) including glutamate and aspartate

This selective pattern—widespread losses in homeostatic connectivity contrasted with gains centred on immunologically relevant pathways—supports the hypothesis that RA involves coordinated metabolic network reorganisation that favours immune activation over normal cellular maintenance [13]. However, these exploratory findings require validation with robust multiple-testing correction, longitudinal data, and functional mechanistic studies before any definitive biological or therapeutic conclusions can be drawn.

### 2.3 Implications for Treatment Stratification

The identification of molecular subtypes independent of ACPA status, combined with selective metabolic network disruption, suggests potential applications for precision stratification in RA, although substantial validation is required. Current treatment recommendations are largely organised by disease activity, prior DMARD exposure, and comorbidities, with serostatus acting primarily as a prognostic modifier rather than a basis for targeted drug selection [25, 26]. At the same time, synovial and transcriptomic studies increasingly show that treatment-relevant heterogeneity is not fully captured by conventional clinical or serological stratification [5, 27].

If validated, molecular profiling could enable several exploratory applications. First, multi-omics signatures might help identify patients more likely to benefit from particular therapeutic classes. For example, patterns dominated by metabolic pathway perturbation could hypothetically favour agents with known immunometabolic effects, such as metformin or small-molecule JAK inhibitors, which already show efficacy in RA and modulate metabolic signalling [28, 29]. Second, molecular subtyping could inform trial design. Phase III RA trials of biologics and targeted synthetic DMARDs often achieve only 40–60% response rates despite standard enrichment strategies, reflecting the pooling of biologically heterogeneous patients [30, 35]. Biomarker-guided enrichment or stratified randomisation based on multi-omics signatures could, in principle, unmask subgroup-specific treatment effects, analogous to biomarker-defined oncology trials such as alpelisib in PIK3CA-mutated breast cancer [31, 32]. Third, network-level metrics—for example, summarising the density or topology of protein–metabolite associations in specific pathways—might serve as quantitative biomarkers that complement conventional composite indices derived from joint counts and acute-phase reactants [33].

**Table 1.** Potential applications and required validation for multi-omics stratification.

| Application | Hypothesis | Validation Needed |
| --- | --- | --- |
| Treatment selection | Subtypes predict drug class response | Prospective trials with pre-treatment profiling |
| Trial stratification | Reduces heterogeneity | Retrospective trial analysis |
| Response monitoring | Network metrics detect residual disease | Longitudinal sampling |

### 2.4 Limitations and Future Directions

This exploratory study has several important limitations. First, the cross-sectional design precludes causal inference: we cannot determine whether the observed molecular subtypes reflect distinct aetiological pathways, transient disease states, or treatment-induced phenotypes, nor whether association network patterns precede treatment response or merely mirror current therapy and disease activity [37]. Second, our network analysis used exploratory correlation thresholds (|*r*| *>* 0.4, *p <* 0.01) without formal control of the false discovery rate. Although permutation testing supported a global difference in edge density (p*<* 0.001) and inflammatory metabolite hyperconnectivity appeared robust across reasonable thresholds, individual edge– and node-level inferences require validation.

Third, medication heterogeneity represents a major uncontrolled confounder. Patients were receiving diverse conventional and biologic DMARDs, and longitudinal multi-omics work in RA has demonstrated that effective treatment can substantially reshape blood-based molecular profiles [38]. Treatment-naïve cohorts and/or pre–post treatment sampling will be essential to disentangle disease-intrinsic signatures from drug effects. Fourth, the modest sample size (80 RA patients, 40 controls) and single-centre recruitment limit statistical power, increase the risk of unstable effect estimates, and constrain generalisability [39].

Future work should prioritise: (1) external validation in independent, multi-centre cohorts with diverse demographics and treatment patterns; (2) prospective studies linking baseline multi-omics profiles and network metrics to subsequent treatment response and long-term outcomes; (3) more rigorous statistical frameworks, including explicit FDR control and threshold-free or weighted network approaches such as WGCNA for robust co-expression and correlation network analysis [40]; (4) mechanistic validation in synovial tissue and relevant immune cell subsets to bridge plasma signatures with site-of-disease biology; and (5) development of analytically robust, cost-effective assays suitable for routine clinical use.

## 3 Conclusion

This exploratory study suggests that integrated proteomic–metabolomic profiling can reveal molecular heterogeneity in RA that is not captured by conventional clinical and serological markers. We addressed two complementary questions: whether RA patients form stable molecular subgroups, and how disease perturbs protein–metabolite association networks.

Consensus clustering identified two reproducible molecular subtypes (overall consensus = 0.898) within the RA cohort, with membership distributed independently of ACPA status. This supports the view that RA comprises multiple pathobiological states not fully reflected by standard serological classification, consistent with synovial and transcriptomic studies showing distinct molecular phenotypes linked to outcome [4, 5, 41].

Network analysis revealed extensive but selective disruption of metabolic coordination in RA. Overall edge density was markedly reduced compared with controls, yet the strength of surviving associations was preserved, indicating targeted pathway disconnection rather than uniform weakening of all relationships. Loss of connectivity was most pronounced in amino-acid metabolism, whereas inflammatory mediators such as glutamate, aspartate and sphingolipids showed relative hyperconnectivity, in line with models of RA as an immunometabolic disease in which immune cells undergo coordinated metabolic reprogramming [19, 21, 42].

Taken together, these findings are consistent with a model in which RA comprises molecularly distinct subgroups characterised by differing patterns of metabolic network reorganisation. In current practice, treat-to-target algorithms achieve clinical responses in only a subset of patients and do not routinely incorporate molecular stratification [1, 34]. If replicated and linked prospectively to treatment outcomes, multi-omics signatures and network metrics could, in principle, support more refined therapeutic selection, inform biomarker-enriched trial designs, and highlight metabolic pathways that complement existing immune-targeted strategies [31, 43].

However, these results remain hypothesis-generating. The cross-sectional design, medication heterogeneity, modest sample size, and exploratory statistical thresholds introduce substantial uncertainty, and we have not yet shown that the identified subgroups or network features predict response to specific therapies [35, 36]. Nonetheless, the convergence of molecular subtyping and network analysis on coherent biological signals—heterogeneity beyond traditional markers and selective metabolic rewiring—supports further investigation of multi-omics approaches as part of an emerging precision-medicine agenda in RA.

